# Multiple-trait subsampling for optimized ancestral trait reconstruction

**DOI:** 10.1101/2022.10.11.511762

**Authors:** Xingguang Li, Nídia S. Trovão, Joel O. Wertheim, Guy Baele, Adriano de Bernardi Schneider

## Abstract

Large datasets along with sampling bias represent a challenge for phylodynamic reconstructions, particularly when the study data are obtained from various heterogeneous sources and/or through convenience sampling. In this study, we evaluate the presence of unbalanced sampled distribution by collection date, location, and risk group of HIV-1 subtype C using a comprehensive subsampling strategy, and assess their impact on the reconstruction of the viral spatial and risk group dynamics using phylogenetic comparative methods. Our study shows that the most suitable dataset for ancestral trait reconstruction can be obtained through subsampling by collection date, location, and risk group, particularly using multigene datasets. We also demonstrate that sampling bias is inflated when considerable information for a given trait is unavailable or of poor quality, as we observed for the risk group in the analysis of HIV-1 subtype C. In conclusion, we suggest that, even if traits are not well recorded, including them deliberately optimizes the representativeness of the original dataset rather than completely excluding them. Therefore, we advise the inclusion of as many traits as possible with the aid of subsampling approaches in order to optimize the dataset for phylodynamic analysis while reducing the computational burden. This will benefit research communities investigating the evolutionary and spatiotemporal patterns of infectious diseases.

## Introduction

Large sequencing efforts have greatly increased the availability of genomic data of infectious agents or pathogens in public databases (1). This data availability has led to the development of novel methods to speed up molecular epidemiological analyses of these datasets (2, 3). Yet, although these tools aim to solve the problem of data processing, they do not by themselves resolve the issue of data representativeness, as seen through the presence of sampling bias in large databases, resulting in datasets with a skewed distribution of certain traits not truly representing the population diversity. Genetic databases, such as GenBank (4) and GISAID (5), are used as repositories for genomic data, which is often deposited at the moment of submission of a manuscript to peerreviewed journals, with a few notable exceptions such as the genomic data be deposited during the Ebola outbreak in West Africa (6), the SARS-CoV-2/COVID-19 pandemic (7, 8) and throughout seasonal influenza virus surveillance efforts to inform vaccine composition (9). In public databases, sampling bias can be seen through the random deposit of samples in the database in an unintended way (sequences being deposited as project-dependent and not population-dependent) that does not reflect a fair representation of the true population, resulting in some traits (i.e., genetic diversity, location, populations at greater risk of HIV acquisition (PGRHA)) of the target population having a lower or higher sampling probability than others compared to their actual prevalence (10–13).

Sampling bias is a persistent concern when performing phylogeographic inference (14, 15). Apart from an increase in taxon sampling having been shown to aid in the reduction of phylogenetic error (16), several software applications target the reduction of size and redundancy for the purpose of phylogenetic analysis (17) or the increase in phylogenetic diversity while reducing data set size (18). Sequencing errors and the lack of a representative sampling from large datasets can lead to inferring erroneous phylogenies, hampering accurate downstream conclusions (19). Sampling bias can, for example, lead to incorrect inference of ancestral locations and migration rates from oversampled regions, leading to spurious results that may affect public policy in the response of an epidemic (14). The presence of sampling bias is challenging for all currently available phylogeographic models, and mitigating such bias might require large data set sizes and the incorporation of associated metadata in those models (20). Additionally, sequencing errors have been shown to affect phylogenetic inference, potentially impacting local SARS-CoV-2 lineage tracing efforts (21).

Subsampling is typically a strategy to mitigate any biases present in a dataset, and thus to improve the representativeness of the actual patterns of the epidemics. However, a recent study has shown that such subsampling strategies do not consistently improve (discrete) phylogeographic inference at intermediate levels of sampling bias, and that the improvements is dependent on the actual migration model (20). Subsampling is also used to reduce the computational burden of phylogenetic and other molecular epidemiology analysis of very large genetic datasets, such as those for the human deficiency virus (HIV) (11), seasonal influenza viruses (22) and the severe acute respiratory syndrome coronavirus 2 (SARS-CoV-2) (23).

Representativeness is multi-dimensional, in the sense that a single genomic dataset does not only consist of genomic data, but multiple underlying metadata layers (traits) which when combined allow for a comprehensive view of the population represented by the dataset. Studies focusing on the spatiotemporal dynamics of pathogens often tend to subsample based on location, particularly for the challenging discrete (location) trait reconstruction analysis (11). However, even when the goal is to purely reconstruct the pathogen’s spatial spread, including more traits during the subsampling process might improve the representativeness of the actual underlying patterns of the epidemics and lead to more accurate results. Among the existing large genomic data repositories, the Los Alamos National Laboratory (LANL) HIV Sequence Database (https://www.hiv.lanl.gov) is one of the most widely used for HIV research. In addition to genomic data, the database contains metadata associated with the viral genetic sequences, including records of the collection date and sampling country along with PGRHA information for certain samples, thus making it an ideal database to evaluate sampling bias on multiple traits associated with the samples. HIV-1 subtypes B and C have the largest number of sequences recorded in the LANL HIV Sequence Database (https://www.hiv.lanl.gov/content/sequence/HIV/mainpage.html). As of August 3 2022, there were 535 995 and 152 290 records for HIV-1 subtypes B and C, respectively. Subtype B is the most widespread HIV-1 variant accounting for approximately 11% of all infections worldwide (24), and has been extensively studied including in the phylodynamic context (25). Despite there being studies addressing the global evolution and spatiotemporal patterns of HIV-1 subtype C (26), to the best of our knowledge the present study is the first one to address the potential influence of sampling bias on the accuracy of such reconstructions.

Many studies tend to focus on genomic analyses for specific regions, however the regional transmission dynamics might not fully represent the overall evolutionary and spatiotemporal patterns of the disease worldwide. In this study, we evaluate the effect of dataset subsampling based on a combination of traits (date, country, PGRHA) on phylogenetic inference and subsequent downstream analysis, such as ancestral trait reconstruction and phylogeographic inference. We subsample and analyze two large HIV-1 subtype C datasets with sequences collected globally obtained from LANL, encompassing near-complete genome and partial *pol* gene, with associated metadata. We find that subsampling using a combination of genetic sequence and metadata traits yields more comprehensive phylogenetic results than the usual subsampling based on a single metadata trait.

## Materials and Methods

### Sequence dataset compilation

All available near-complete genome sequences (HXB2 genome position 790–9417, with minimum fragment length of 6 000 bp) and partial *pol* sequences (HXB2 genome positions 2 200–3 500, with minimum fragment length of 600 bp) of HIV-1 subtype C with known sampling dates and geographic information were retrieved from the Los Alamos National Laboratory (LANL) HIV Sequence Database (https://www.hiv.lanl.gov) on March 26th, 2021. Problematic sequences, as defined by LANL, were removed and only one sequence per patient was selected before download. Sequence quality was analyzed using the Quality Control tool from the LANL site, and all genotype assignments were confirmed using RIP v.3.0 (27). Hypermutation analysis was performed using Hypermut v2.0 (28). The two final datasets include 1 221 publicly available near-complete genome sequences of HIV-1 subtype C (*full1221*) with known sampling year (1986–2019) and locations (32 countries), and 34 229 publicly available partial *pol* sequences of HIV-1 subtype C (*pol34229*) with known sampling year (1986–2019) and locations (106 countries). For both *full1221* and *pol34229* datasets, we grouped PGRHA into six categories: male who have sex with male (SM), people who inject drugs (PI), heterosexual (SH), mother-to-baby (MB), not recorded (NR), and other (OT), as described at LANL (https://www.hiv.lanl.gov/content/sequence/HIV/data_dictionary/data_dictionary.html).

### Molecular sequence analyses

The *full1221* and *pol34229* datasets were processed separately. Multiple sequence alignments of the two datasets (*full1221* and *pol34229*) were obtained using MAFFT v7.427 (29) under an automatic algorithm and subsequently adjusted manually in BioEdit v7.2.5 (30). Next, we excluded sequences with more than 50% gaps as well as duplicate sequences, defined as having the same collection date, country, PGRHA, and nucleotide sequence. This resulted in a full genome dataset of 1 210 sequences (*full*), and a pol gene dataset comprising 33 859 sequences (*pol*).

In order to subsample the datasets, we used the Subsampling According to Metadata for Phylogenetic Inference (SAMPI) python tool (available at: https://github.com/jlcherry/SAMPI). Subsampling was performed to obtain a homogeneous collection of samples using the variables country, PGRHA, and year while maintaining a manageable dataset size lower than 1000 sequences for computational efficiency. Three subsets with repetitions for full genomes and pol gene were assembled. (CP) country, PGRHA, and year, (C) country and year, and (P) PGRHA and year. This resulted in the following datasets: (fullCP): 10 sequences per date, country, and PGRHA, n=626 sequences; (fullC): 10 sequences per date and country, n=562 sequences; (fullP): 10 sequences per date and PGRHA, n=393 sequences; (pol CP): 1 sequence per date, country, and PGRHA, n=698 sequences; (pol C): 1 sequence per date and country, n=986 sequences; (pol P): 7 sequences per date and PGRHA, n=727 sequences. We selected a higher number of sequences per date and PGRHA for pol C given the smaller number states of PGRHA and with the objective of having a subsampled dataset of similar size to the other datasets.

To examine the reproducibility of the datasets and analysis, we performed three independent repetitions of each subsampling strategy. Multiple iterations of maximum-likelihood (ML) phylogenetic reconstruction using RAxML v8.2.12 (31) under a GTR+Γ_4_+I nucleotide substitution model with 1 000 bootstrap replicates were performed, with removal of outlier sequences – those with incongruent sampling dates and root-to-tip genetic divergence – via the TempEst software package v1.5.3 (32). This resulted in full genome datasets with 626 (*fullCP*), 562 (*fullC*), and 393 (*fullP*) sequences, and partial pol gene datasets with 986 (*pol CP*), 698 (*pol C*), 727 (*pol P*) sequences.

### Phylogenetic reconstruction

ML phylogenetic recon-struction was performed for the original datasets and their subsampling replicates (*full, fullCP, fullC, fullP, pol CP, pol C*, and *pol P*) using RAxML v8.2.12 (31) under the GTR+Γ_4_+I nucleotide substitution model with 1,000 bootstrap replicates. Due to the large size of the *pol* dataset, ML phylogeny reconstruction was performed using IQ-TREE v2.1.2 with the GTR+F+R10 substitution model (33). In addition, we used FigTree v1.4.4 (34) to visualize and annotate the phylogenetic trees with geographic location and PGRHA.

### Phylogenetic tree comparison

In order to understand how subsampling affected the tree topology among the shared taxa among all trees for pol and for the full genome datasets, we compared the tree topologies using the ClusteringInfoDist metric (see below), which provides a similarity score between trees with the same sequences as tips. To this end, we first extracted subtrees from each dataset containing the intersect of the taxa present in all the trees using *ybyra_pruner*.*py* from the YBYRÁ package (35). Then, we separately compared each subtree set from the near-complete genome and partial pol gene using the “ClusteringInfoDist” function from the TreeDist package as implemented in R (36– 38). The ClusteringInfoDist algorithm calculates a normalized tree similarity and distance measures based on the amount of phylogenetic or clustering information that two trees hold in common, where a lower value corresponds to trees that are topologically more similar, with a zero distance corresponding to identical trees. The normalization process on ClusteringInfoDist allows for a better comparison between the results of analyses coming from distinct datasets (i.e, results from pol comparisons vs results from full comparisons). We calculated the average, mean, and performed a two-tailed distribution t-test assuming two-samples of unequal variance (heteroscedastic) in order to identify statistical significance (<0.01) between the ClusterInfoDist values for the groups of subsampled trees.

### Transmission Networks

To evaluate the robustness of our subsampling method, we generated transmission networks based on our phylogenetic reconstructions for all the datasets using the parsimony ancestral reconstruction method in StrainHub v1.1.2. Disease transmission networks of traits of interest display the connectivity and dynamics within each trait (i.e., country and PGRHA) and allow us to understand the behavior the disease and importance of each network node on the spread of the disease (e.g., if there is a single node in the network being a super spreader or if the spread is balanced among nodes).

StrainHub generates a transmission network based on character state changes in metadata, such as collection location, mapped on the phylogenetic tree. The nodes of this transmission network represent the relationship of the ancestral and descendant states of the pathogen sequences (e.g. changes in geography, host shifts, and among PGRHA) (39). We evaluated to what extent subsampling interfered with the structure of the networks by comparing the networks directly and through the centrality metrics of each network (40). Metadata were extracted from the sequence headers and geographic coordinates were extracted from latlong.net. We ranked the datasets’ metadata (country and PGRHA) by degree centrality and Source Hub Ratio (SHR). Degree centrality is defined as the number of edges a trait state has within the network, meaning that the higher the degree, the more connected the state is to other states. The estimates associated with SHR, a score that ranges from 0 to 1, indicate a sink or source behaviour of a particular state, respectively (hub has a SHR=0.5), as implemented in StrainHub (39). We also calculated the Pearson product-moment correlation coefficient for all the pairs of trait states for all original and subsampled *full* and *pol* datasets to understand how subsampling affects the overall transmission network structure.

## Results

### Subsampling

In the *full* dataset, genome sequences collected in South Africa (ZA; 49.7% (601/1210)) and Zambia (ZM; 8.1% (219/1210)), and from the NR (68.6% (830/1210)) and SH (26.9% (325/1210)) populations at greater risk of HIV acquisition (PGRHA) are over-represented compared to the numbers for the other countries and PGRHA (Figure 1). Our subsampling strategy resulted in datasets with the following reduced sequence counts (average between three repetitions of subsampled datasets) for South Africa and Zambia: 31.2% (195/626) and 14.4% (90/626) in the *fullCP* datasets, 26.9% (151/562) and 13.5% (76/562) in the *fullC* datasets, and 34.9% (137.3/393) and 14.0% (55/393) in the *fullP* datasets. Similarly, the sub-sampling by PGRHA results in datasets with the following NR and SH genome sequence counts: 60.9% (381/626) and 31.5% (197/626) in the *fullCP* datasets, 62.1% (349/562) and 30.2% (170/562) in the *fullC* datasets, and 54.7% (215/393) and 33.1% (130/393) in the *fullP* datasets, respectively.

**Fig. 1.**
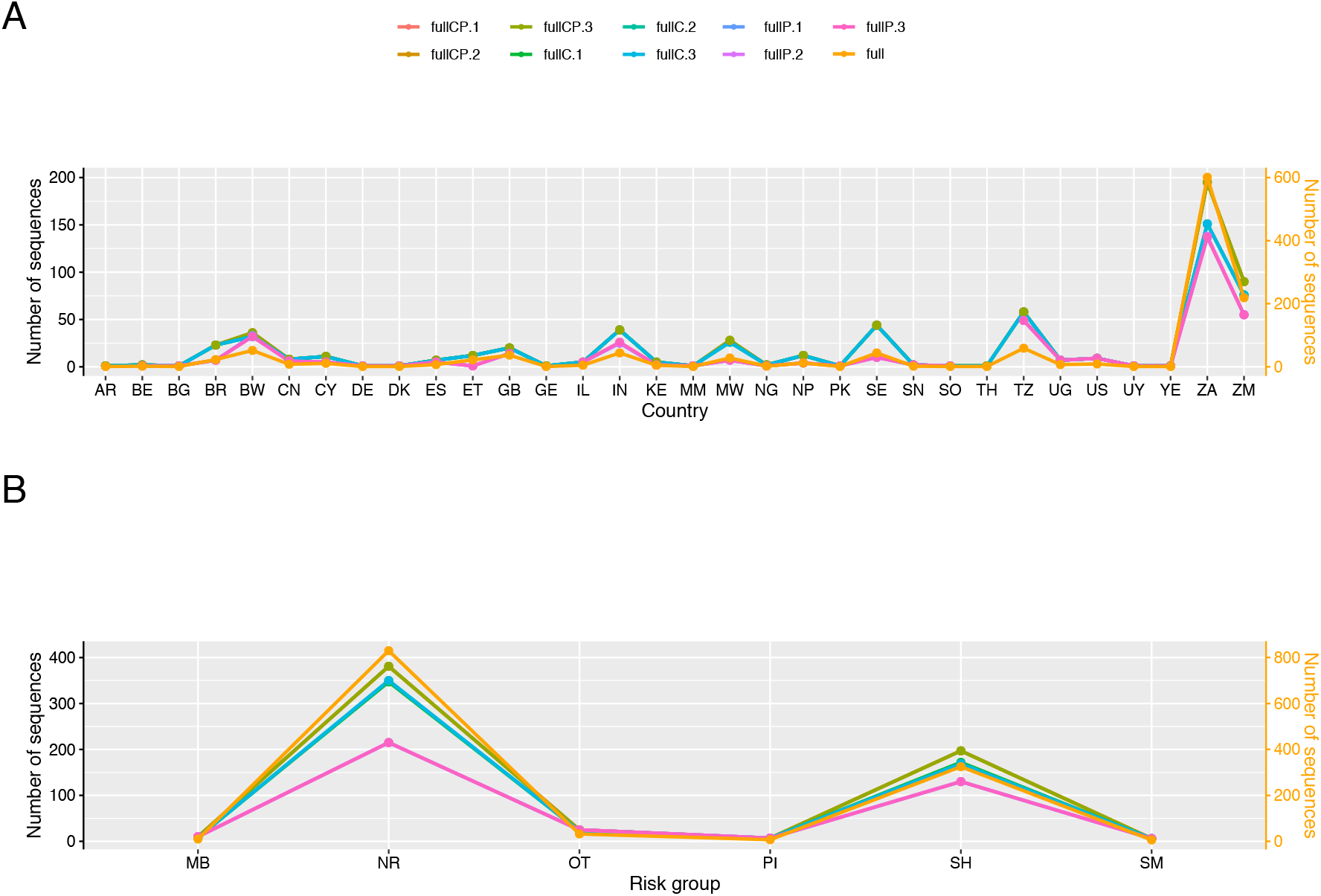
Sampling distributions of metadata traits for the full and subsampled datasets of HIV-1 subtype C. (A) Country distribution for the full and subsampled datasets. The distribution of the original dataset shows a disproportionate amount of samples sampled from BR, BW, IN, MW, SE, TZ, ZA and ZM. (B) Populations at greater risk of HIV acquisition (PGRHA) distribution for the full and subsampled datasets. The distribution of the data shows a large amount of missing data (labeled NR) and higher amount of SH in comparison to the other PGRHA. The number of sequences for the full dataset is labelled on the right y-axis.

In the *pol* datasets (Figure 2), the partial *pol* gene sequences collected in South Africa (51.1% (17312/33859)) and India (8.6% (2922/33859)), and from the NR PGRHA (90.4% (30615/33859)) are over-represented. After subsampling, the average between three repetitions of subsampled datasets for the partial *pol* gene sequences obtained in South Africa and India account for 6.8% (67/986) and 5.7% (56/986) in the *pol CP* datasets, 4.0% (28/698) and 3.3% (23/698) in the *pol C* datasets, and 24.4% (177.3/727) and 13.9% (101.3/727) in the *pol P* datasets, respectively. Likewise, the subsampled partial *pol* gene sequences collected from the NR PGRHA now account for 62.7% (618/986) from the *pol CP* datasets, 75.2% (524.7/698) from the *pol C* datasets, and 26.5% (193/727) from the *pol P* datasets.

**Fig. 2.**
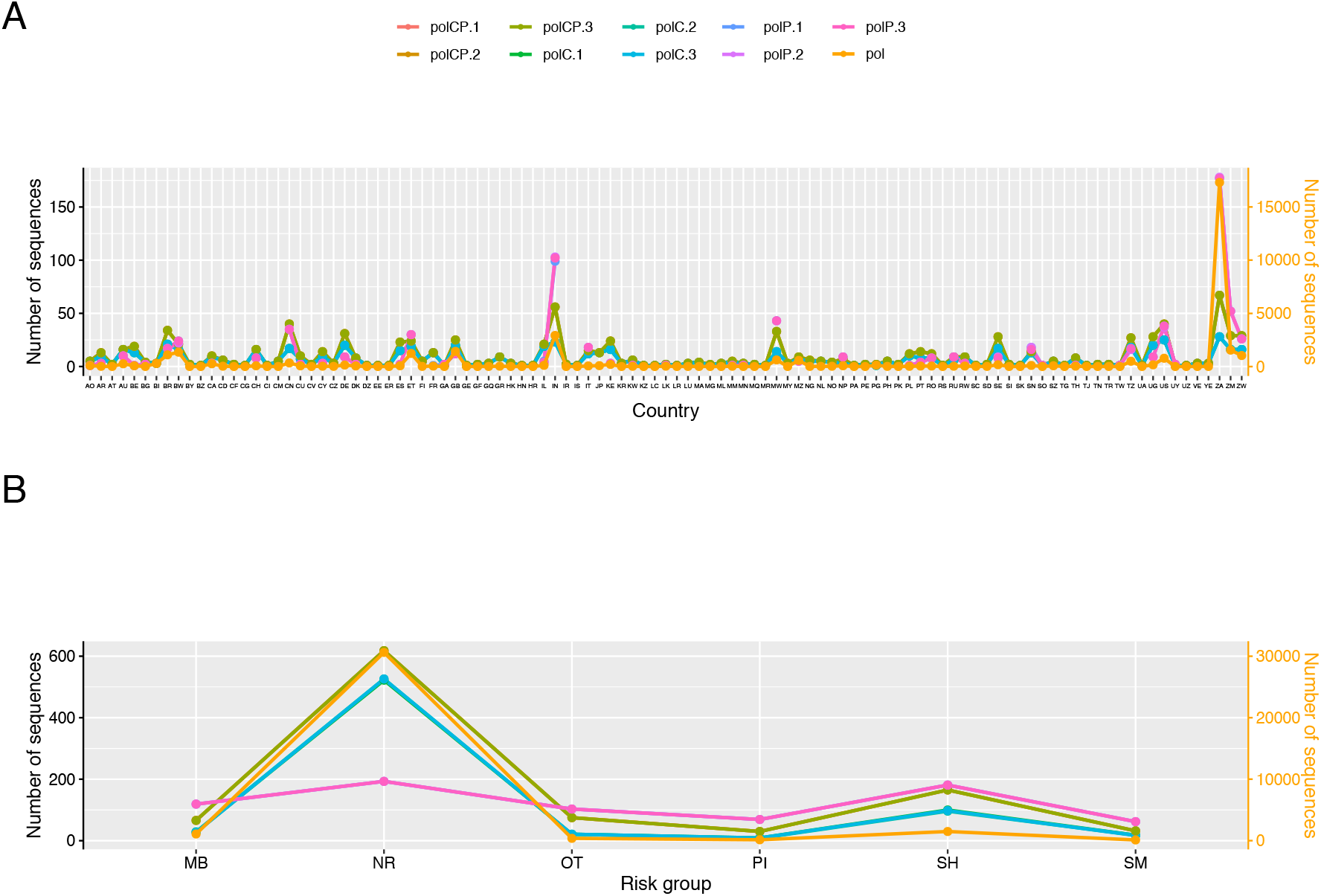
Sampling distributions of metadata traits for the pol and subsampled datasets of HIV-1 subtype C. (A) Country distribution for the pol and subsampled datasets. The distribution of the original dataset shows a larger amount of samples sampled from BR, BW, ET, GB, IN, MW, MZ, TZ, US, ZM and ZW, with a disproportionate amount of samples from ZA. (B) Populations at greater risk of HIV acquisition (PGRHA) distribution for the pol and subsampled datasets. The distribution of the data shows a large amount of missing data (labeled NR) and slightly higher amount of SH in comparison to the other PGRHA. The number of sequences for the pol dataset is labelled on the right y-axis.

The subsampling of the pol P datasets yield a different country composition in comparison to the other datasets (46 or 47 of 105 countries), given the large amount of data and the subsampling method that did not include country as a sub-sampling trait. For downstream analyses, we compared only the intersect of data between each dataset, i.e, pol (105 countries) vs pol CP (105 countries); pol CP (105 countries) vs pol C (105 countries); pol, pol CP, or pol C (46 or 47 of 105 countries) vs pol P (46 or 47 countries).

### Tree comparisons

For the full datasets (Figure 3A), the topologies of the subtrees subsampled by country (fullC; average = 26.24) are the closest in similarity to that of the original dataset (full). Nevertheless, fullCP datasets (average = 28.06) have very close values to fullC, with fullP (average = 33.94) being the most distant datasets to full. For the pol gene dataset (Figure 3B) the topologies of all the subsampled subtrees are mostly equidistant to the original pol dataset (pol CP average = 34.92; pol C average = 34.87; pol P average = 36.02). We estimated that the datasets were not significantly different across full and pol subsamplings, with the exception of fullCP vs fullP (p-value = 0.009) and fullC vs fullP (p-value = 0.0005). Nevertheless, we observe overall similar values across subsamplings for both the full and pol datasets (small variance across subsamplings; fullCP = 2.47; fullC = 0.72; fullP = 0.90; pol CP = 1.10; pol C = 2.49; and pol P = 3.57).

**Fig. 3.**
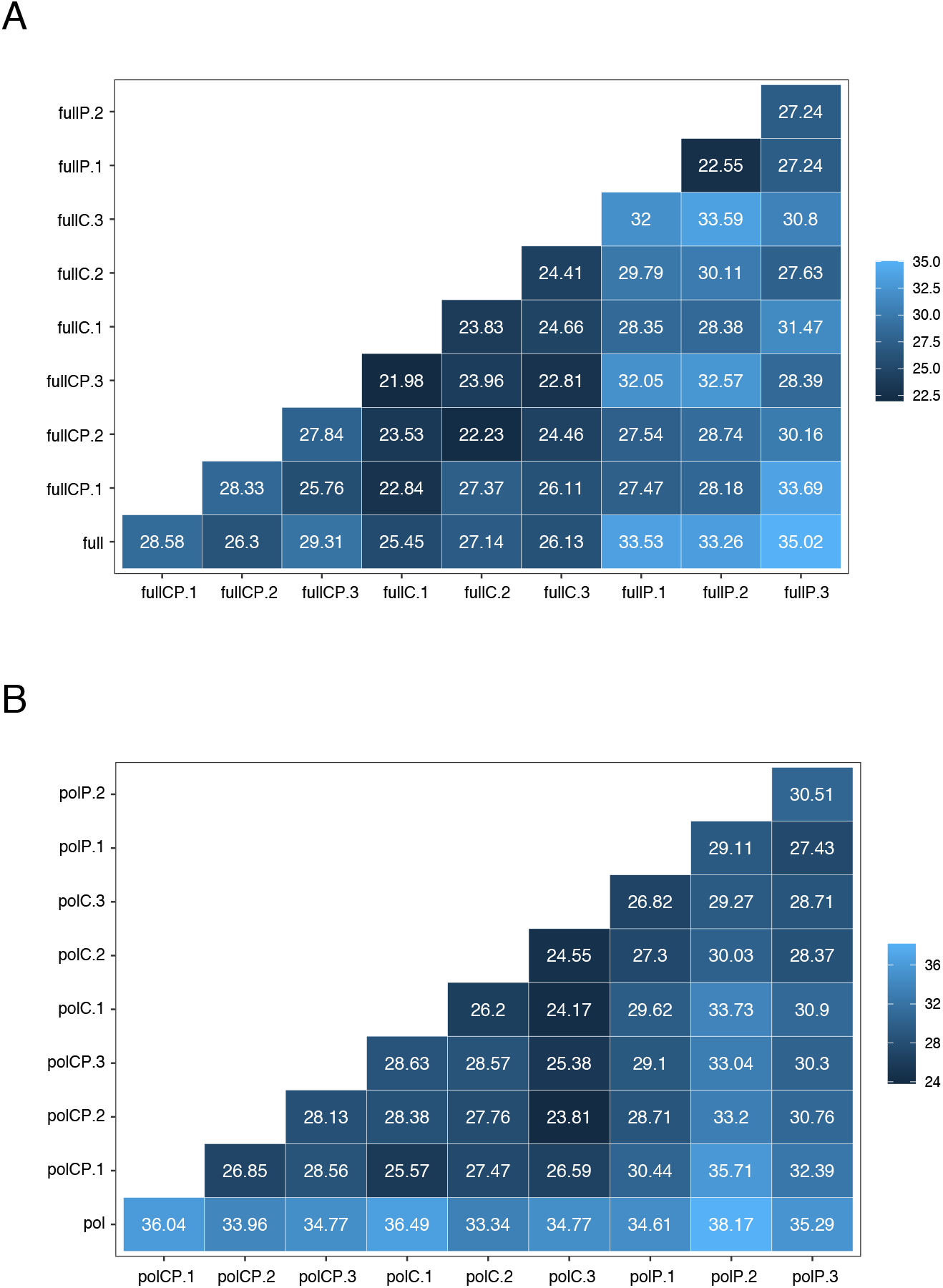
Cluster Info Distance comparison of the phylogenetic topologies of the *full* (A) and *pol* (B) and respective subsampled datasets subtrees. Zero cluster info distance equals identical trees. The topologies of the full and pol subsampled dataset subtrees are overall similar to that of their respective original datasets.

### Transmission networks

We generated transmission networks for all the full and pol datasets. We observed that there was limited variation of the correlation of the degree centrality metric between country and PGRHA with the original dataset across repetitions of the same subsampling strategy or across the three subsampling strategies for full and pol datasets, with the exception of pol P (Figure 4, Supplementary Figures S1, and S2). This means that the degree of connectivity of each country or PGRHA node in the overall transmission network is maintained irrespective of the sub-sampling strategy. Despite the overall maintenance of the country and PGRHA node importance, their behaviours (i.e, sink/hub or source of disease), assessed using the Source-Hub Ratio (SHR) estimate, varies with the subsampling strategy employed. We observed that the correlation of the SHR with the original dataset for the country trait is highest for fullCP and fullC, and lowest for fullP. This pattern is also observed for the pol subsamplings, even though the overall SHR correlations with the original dataset are lower than those for the full dataset.

**Fig. 4.**
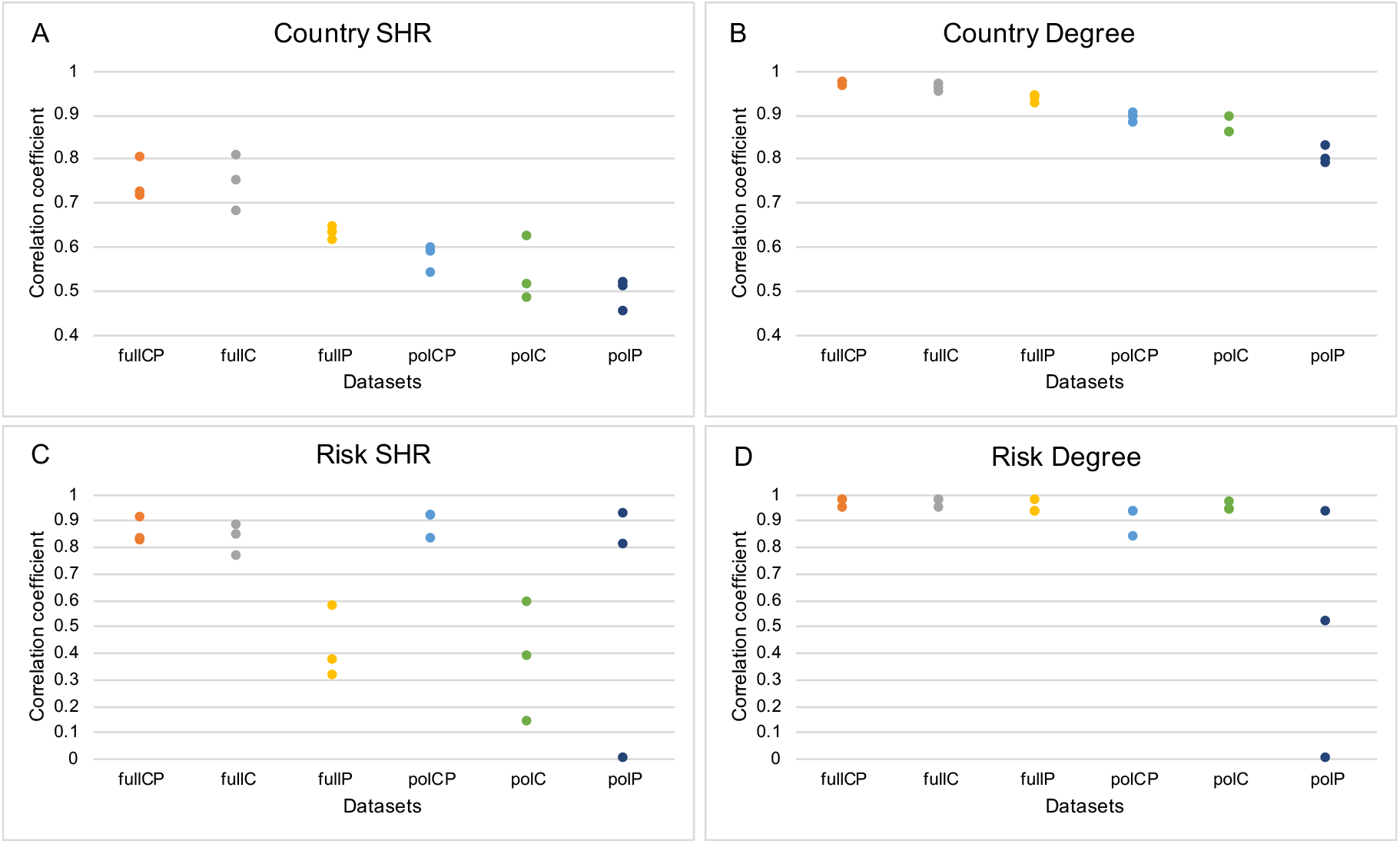
Correlation of Source Hub Ratio (SHR) and degree centrality metrics as a proxy for transmission network structures of HIV-1 subtype C for the *full* and *pol* with respective subsampled datasets by date, country, and populations at greater risk of HIV acquisition (PGRHA). The correlation of the transmission network is higher (thus, similar structures) as the correlation coefficient approximates to 1. (A) Estimates of similarity of the spatial transmission network structure for all subsampled datasets based on SHR metric; (B) Estimates of similarity of the spatial transmission network structure of HIV-1 subtype C for all subsampled datasets based on the degree centrality metric; (C) Estimates of similarity of the PGRHA transmission network structure of HIV-1 subtype C all subsampled datasets based on SHR metric; (D) Estimates of similarity of the PGRHA transmission network structure of HIV-1 subtype C for all subsampled datasets based on the degree centrality metric. The degree of connectivity of each country or PGRHA node in the overall transmission network is generally maintained irrespective of the subsampling strategy, however there is a discrepancy of the country and PGRHA node behaviours as indicated by the varying SHR per subsampling strategy.

Interestingly, the correlation of SHR for the PGRHA trait is higher for both fullCP and fullC than for fullP. This might suggest that for the full dataset, even when subsampling is done solely using country, we obtain a distribution of samples such that they are similar to the overall PGRHA trait structure of the original full dataset (Figure 5). However, the limited information for PGRHA yields results that do not represent the overall PGRHA trait structure in the original dataset. This behaviour is not observed in the pol subsampling, where pol CP and pol P have the highest SHR correlation with the original dataset, agreeable with the fact that both strategies include information for PGRHA, whereas pol C yields the lowest correlation with the original PGRHA transmission network.

**Fig. 5.**
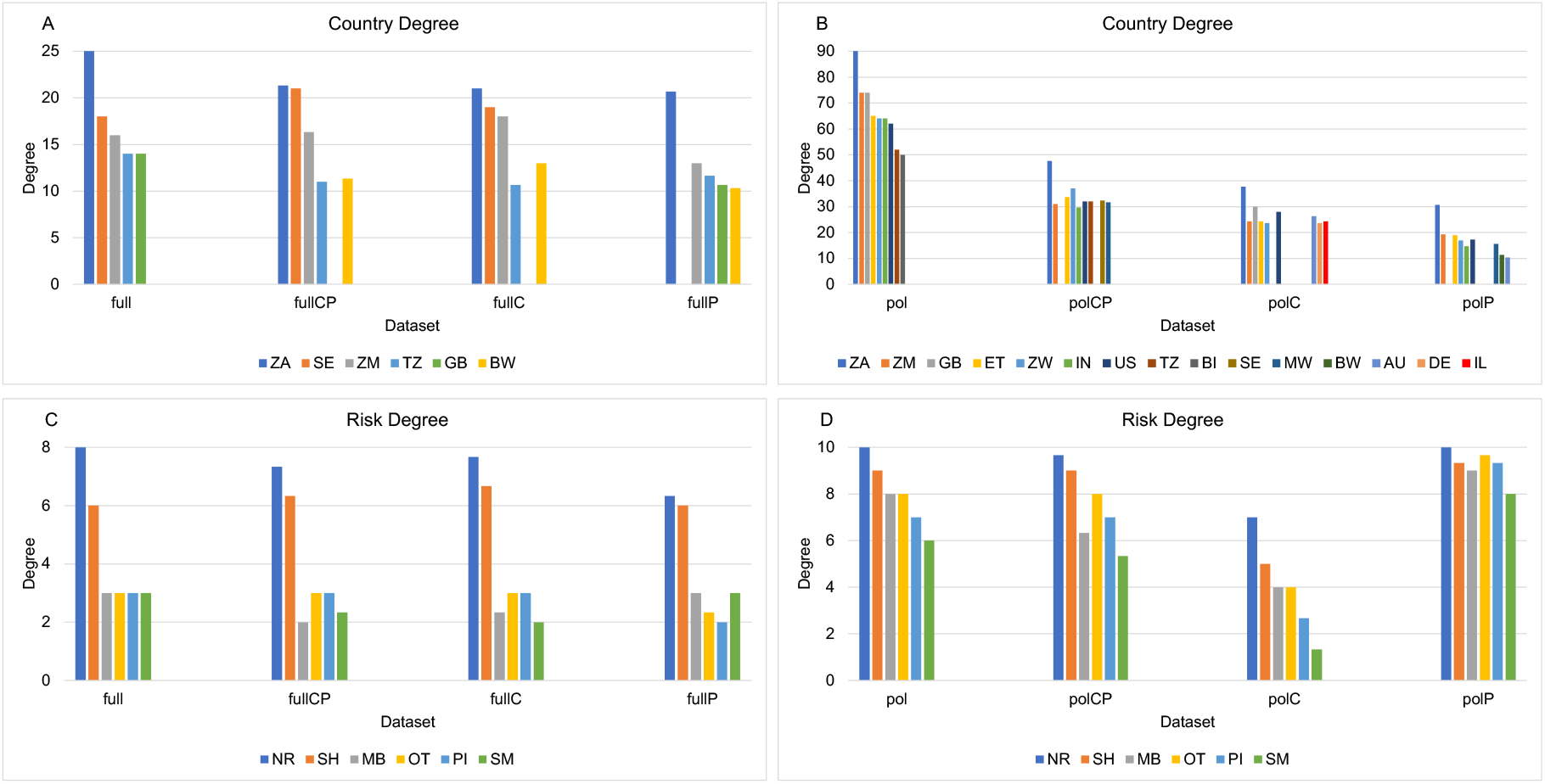
Distribution of the average degree centrality among all subsamples for the top clusters in the pol and full subsets. (A) Degree centrality for top five countries (country trait) for full subsets; (B) Degree centrality for top nine countries (country trait) for pol subsets; (C) Degree centrality for all populations at greater risk of HIV acquisition (PGRHA) for full subsets; (D) Degree centrality for all PGRHA trait for pol subsets. Including country and PGRHA as subsampling traits yields the most consistent results for both traits’ transmission networks. The distribution of degree centrality among the nodes of the networks evaluated show that datasets subsampled by PGRHA and country or solely by country result in patterns similar to those for the original pol and full datasets. The top countries in terms of degree centrality are mostly conserved across the full datasets, with wider variance observed in the pol datasets likely due to the larger number of locations for the country trait in these.

Looking within the transmission networks geographically, in the full genome dataset, the countries with the highest degree centrality are (in order of high to low degree centrality) South Africa (ZA), Sweden (SE), Zambia (ZM), Tanzania (TZ) and the United Kingdom (GB) (Figure 5A). However, GB does not rank among the top five countries in the fullCP and fullC datasets, where it is replaced by Botswana (BW). Moreover, Sweden does not rank among the top five countries in the *fullP* dataset, but it does include GB and BW. In the pol datasets, given the larger number of countries, we elected to display the top nine countries for each dataset. The countries with the highest degree centrality are (in order of high to low degree centrality) ZA, ZM, GB, Ethiopia (ET), Zimbabwe (ZW), India (IN), United States (US), TZ and Burundi (BI). The polA datasets best represent the pol dataset’s top 10, with only two countries being replaced (GB and BI by SE and Malawi; MW), with these two countries being replaced as numbers 8 and 9. The polB dataset replaced three of the top 9 countries from the original dataset (IN, TZ, BI with Australia (AU), Germany (DE) and Israel(IL)), and polC replaced three countries as well (GB, TZ, BI with AU, BW, MW) (Figure 5B).

We have also summarized the spatial transmission dynamics among the original full and pol datasets and their respective subsampled dataset. For the full datasets (Supplementary Figure S3), even though there was some variation across replicates, we observed an overall similar pattern characterized by viral dissemination from Africa to Europe and viceversa, as well as from Africa do the US and South Asia. The pol dataset (Supplementary Figure S4) depicts more complex spatio-temporal dynamics that included the viral movements described for the full datasets, plus introductions from North America to Asia, Europe to South America, and Africa to Oceania. The patterns observed for pol C were more sparse, likely due to the reduced number of countries included for this dataset compared to the other pol datasets.

The transmission network among PGRHA shares a similar result across all of the full datasets, with SH and NR having the highest mean degree centrality, therefore contributing with the highest number of connections within the network, and OT, PI, MB and SM contributing less (Figure 5C). For the pol datasets, we see similar results with NR and SH having the highest for pol, polA and polB, and a more elevated degree centrality for OT, PI, MB and SM, with polC having a high degree centrality on all PGRHA with an almost uniform distribution (Figure 5D).

The summarized PGRHA transmission dynamics among the original full and pol datasets and their respective subsampled dataset highlight the results described in the paragraph above. In both the full and pol datasets (Supplementary Figures S5 and S6), we reconstructed similar PGRHA dynamics with NR acting as the main source among PGRHA followed by the SH group, irrespective of subsampling. In the full datasets, we estimated the large majority of transmission dynamics occurring from NR to SH. In the pol datasets, the viral transmissions were mostly from NR to SH but we also estimated a substantial proportion of viral seeding from NR to MB. The patterns observed for pol C were unlike those observed for any other dataset, with no particular PGRHA standing out in terms of viral source or sink. This is again likely due to the reduced number of countries included in this dataset compared to the other pol datasets.

## Discussion

In this study, we investigated HIV-1 subtype C evolutionary and spatiotemporal dynamics while subsampling the genetic data to decrease the sequence counts from over-sampled traits. Subsampling was performed in order to mitigate biases introduced during sampling of PGRHA, as well as of countries through time. To this end, we compiled comprehensive sequence datasets of full genomes and the pol gene region, and revealed that both datasets contained inherent biases irrespective of the trait studied, as observed through the heterogeneous distribution of the datasets (Figure 1 and 2). We could not compare the subsampled dataset distributions to the real population case estimates, which are impossible to obtain. The available epidemiological curves are not desegregated by HIV-1 type and are biased by time and spatial surveillance coverage and effectiveness (41). For these reasons, we assumed for this study that an unbiased sampling should follow a near uniform distribution.

Sampling strategies and procedures like subsampling methods can help address varying trait representativeness in the metadata associated to genomic datasets. There are many studies that apply subsampling methods in an attempt to correct for bias on sampling date and location. For instance, studies targeting the early spread and epidemic ignition of HIV-1 in humans (11), investigating the spatial history of HIV-1 subtype B in the United States (25), and exploring the rapid epidemic expansion of the SARS-CoV-2 Omicron variant in southern Africa (13). Nevertheless, these studies do not comprehensively examine the effects of varying representativeness of traits and its implications on phylodynamic reconstruction. Here, we have observed that a more comprehensive subsampling strategy that includes as many traits as possible (date, location, PGRHA) yields the best result in retaining the original dataset properties, as demonstrated by the high similarities of the transmission networks between the HIV-1 subtype C full and pol, and the fullCP and polCP datasets, respectively (Figure 4). Furthermore, studies that take into account sampling bias are often limited to a single replicate of a particular subsampling method (11, 13, 42, 43). We have demonstrated that this likely does not have harmful implications to the interpretation of the results as there is little variation of the overall tree topology across subsampling replicates (Figure 3), as well as in the ancestral trait reconstruction (Figure 4).

Comparing the tree topologies of the original full and pol datasets with their respective subsampled datasets allows uncovering which subsampling strategies best represent the original structure and whether that structure is punctuated by a particular trait. Our analyses indicate that both full and pol along with their subsampled datasets present comparable variability in tree topology among the subsets both in terms of country and PGRHA. However, there are inherent limitations in both datasets as observed by the majority of sequences labeled as NR (PGRHA) in all datasets.

Our analyses indicate that there is a slightly stronger signal in the full dataset for location as shown by the smaller distances across the original dataset and those subsampled using the location trait, whereas the pol dataset seems to hold the same level of information for both country and PGRHA traits, indicating a more balanced dataset. The comparable ClusterInfoDist metric across datasets and respective subsample repetitions suggest that irrespective of the subsampling strategy the overall structure of the original topology is maintained given the similar values across all comparisons (Figure 3). We can assume that the reason behind the original full and pol datasets having a slightly lower degree of similarity, as measured through ClusterInfoDist, to the datasets subsampled by PGRHA (fullP and polP) might be that the datasets are mainly driven by location, regardless of the skewed sampling distribution in certain geographical locations, such as India and South Africa (Figures 1 and 2), which may be in part due to the predominance of the HIV-1 subtype C in these regions (44). In this scenario, subsampling solely by PGRHA would have a stronger effect on the tree topology, possibly due to the different behavior of the PGRHA within each country. In addition to that, subsampling by PGRHA produces a distinct outcome from subsampling by country, or by both country and PGRHA, most likely owing to the large presence of missing data labeled as NR.

Consequently, the limited information for PGRHA in both full and pol datasets produces inconsistent transmission networks. Overall, these results indicate that the full dataset might be a better choice to investigate phylodynamic patterns since the subsampled datasets produce transmission networks that have higher correlations to the original dataset for both country and PGRHA, and yield more comprehensive evolutionary histories. This result demonstrates once more how multigene datasets provide higher accuracy in phylogenetic analysis despite lower dataset sizes (45).

Geographically, the top countries for the spread of HIV-1 subtype C, as measured by degree centrality of the nodes within the transmission network on both full and pol datasets, are in line with previous studies (44). The most prominent PGRHA in both original full and pol networks is SH, as seen by the larger sampling of heterosexual individuals in these datasets (Figures 1 and 2), which likely represents the current state of the HIV-1 subtype C epidemic at global scale (46). In pol, we also observed MB as a major PGRHA acting as a transmission source. NR seems to be largely connected to SH in both datasets, indicating that the vast majority of non-reported PGRHA may belong to SH (Supplementary figure S5 and S6). The behavior of PGRHA is expected to be dependent on regional norms (47–49), thus the lack of coverage of locations in polP may be the reason why these results diverge considerably from those of the other datasets. Additionally, the countries excluded from polP might be those that have a stronger signal for the dynamics observed in the other datasets, namely transmission events from NR to SH and from NR to MB. Therefore, as expected, including both country and PGRHA as subsampling traits yields the most consistent results for both country and PGRHA transmission networks.

Our attempts to mitigate bias by employing multiple sub-sampling strategies are not without limitations. For instance, since they rely on the metadata available for the genetic data, we might not address biases created by unknown factors. In this HIV-1 subtype C study, most of the metadata regarding PGRHA is not recorded, and NR accounted for 68.6% and 90.4% of sequences in the full and pol datasets, respectively. Besides, some of the metadata may be mislabeled, such as reports of “men who have sex with men” (SM) due to HIV/AIDS-related stigmatization and discrimination, as reported in previous studies (50). Both sequence data and associated metadata are critical to gain more detailed insights into the evolutionary and spatiotemporal patterns of HIV-1 subtype C and other pathogens. Therefore, more reporting and sharing of data in an open and real-time fashion is needed for an effective public health response.

Comparing the original and subsampled datasets to epidemiological data could be a solution to the present issue in sampling. However, this kind of data also often suffers from biases, including those created by under-sampling in low- and middle-income countries or are not documented particularly for the early dynamics of the epidemics (51, 52). We made an effort to obtain retrospective epidemiological data documenting the number of patients infected with HIV-1 subtype C, but the epidemiological data does not report nor is sorted by subtype, which further complicated this endeavour. Additionally, HIV/AIDS being a disease associated with severe stigma could lead to case reports that do not accurately represent the overall circulation patterns (50, 53).

Even though we here offer a detailed approach to reduce inherent biases and further optimize ancestral trait reconstruction by subsampling large datasets, there are other procedures to account for issues with sampling, including careful research and surveillance design, simulations, and weighted methods based on metrics such as prevalence (10, 54–58). New methodological developments enable phylogeographic inferences that are not as affected by sampling bias (14), but currently do not scale well with the increasing number of sequences and locations, and hence make analysis of large data sets computationally challenging.

In summary, we address the challenges of working with large datasets and sampling bias using a subsampling approach based on date, country, and PGRHA. We evaluate how this approach can mitigate bias and optimize data analyses based on the available metadata. We also highlight the importance of rigorously recording metadata in addition to the genetic sequences. This study systematically evaluates strategies to optimize ancestral trait reconstruction in HIV-1 subtype C, and will be helpful to future phylodynamic analysis of this virus, as well as serve as a reference to the study of other pathogens.

## Acknowledgements

We would like to thank Joshua L. Cherry from the National Center for Biotechnology Information, National Library of Medicine, National Institutes of Health, Bethesda, Maryland, USA for his contributions to the subsampling script used in this study. We also would like to thank François Cholette from the National HIV and Retrovirology Laboratories, National Microbiology Laboratory at the JC Wilt Infectious Disease Research Centre, Public Health Agency of Canada, Winnipeg, Canada for the insightful discussions during the development of this manuscript. The opinions expressed in this article are those of the authors and do not reflect the view of the National Institutes of Health, the Department of Health and Human Services, or the United States government.

