## Supplementary figures for "Multiple-trait subsampling for optimized ancestral trait reconstruction"

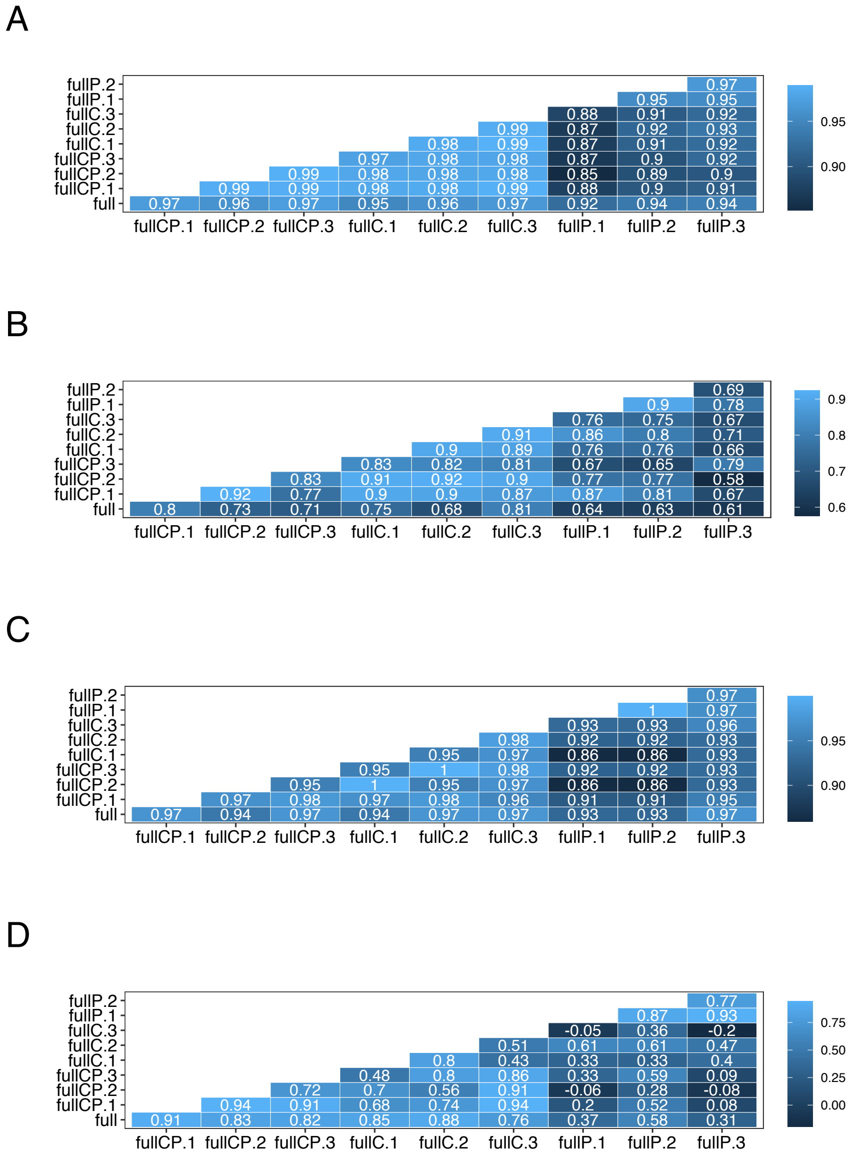


Figure 1: Correlation of network centrality metrics for complete genomes (full). A: Country trait / degree centrality, B: Country trait / Source Hub Ratio, C: PGRHA trait / degree centrality, D: PGRHA trait / Source Hub Ratio.

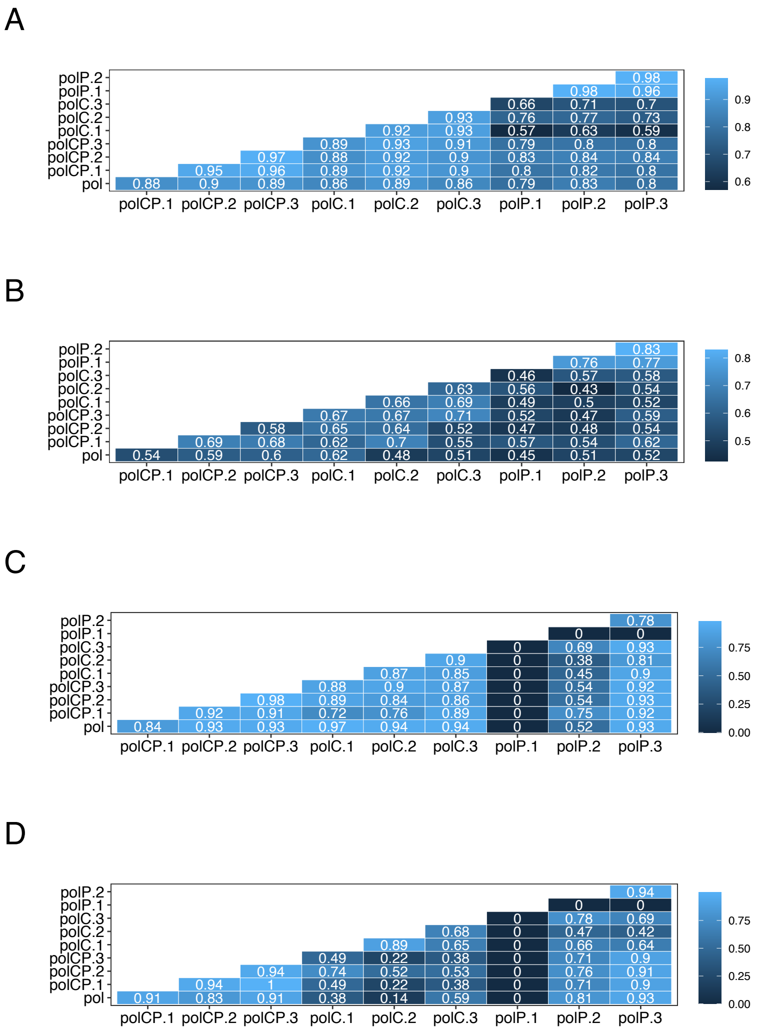


Figure 2: Correlation of network centrality metrics for the pol gene. A: Country trait / degree centrality, B: Country trait / Source Hub Ratio, C: PGRHA trait / degree centrality, D: PGRHA trait / Source Hub Ratio.

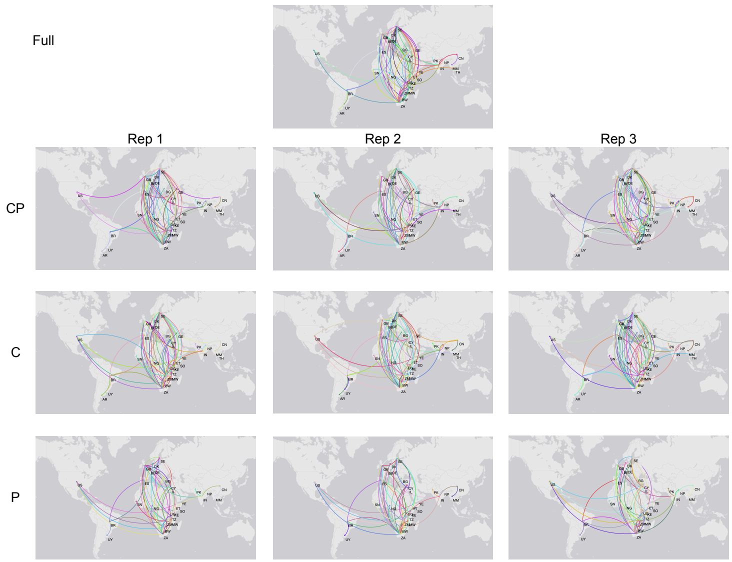


Figure 3: Reconstruction of HIV-1 subtype C global spatial transmission network using the *full* and subsampled datasets visualized using StrainHub v.1.1.2.

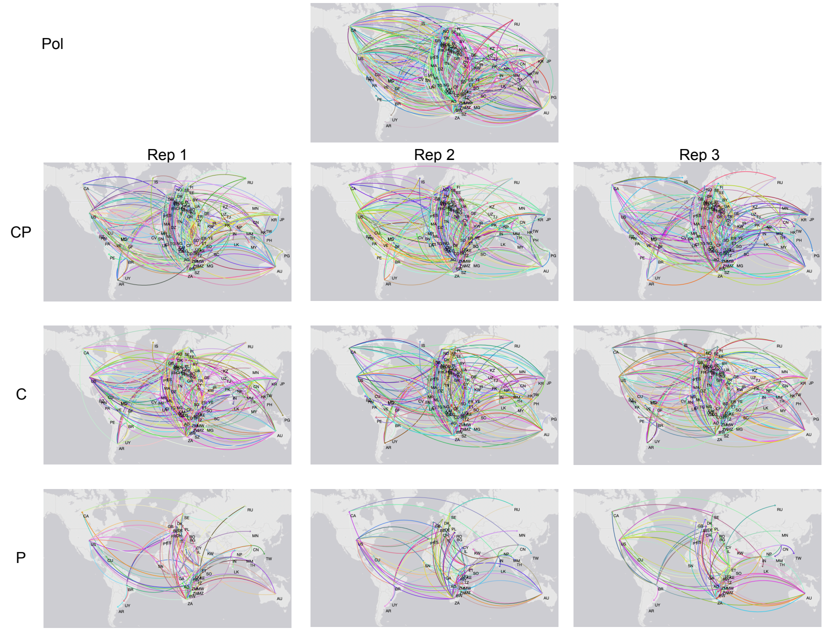


Figure 4: Reconstruction of HIV-1 subtype C global spatial transmission network using the *pol* and subsampled datasets visualized using StrainHub v.1.1.2.

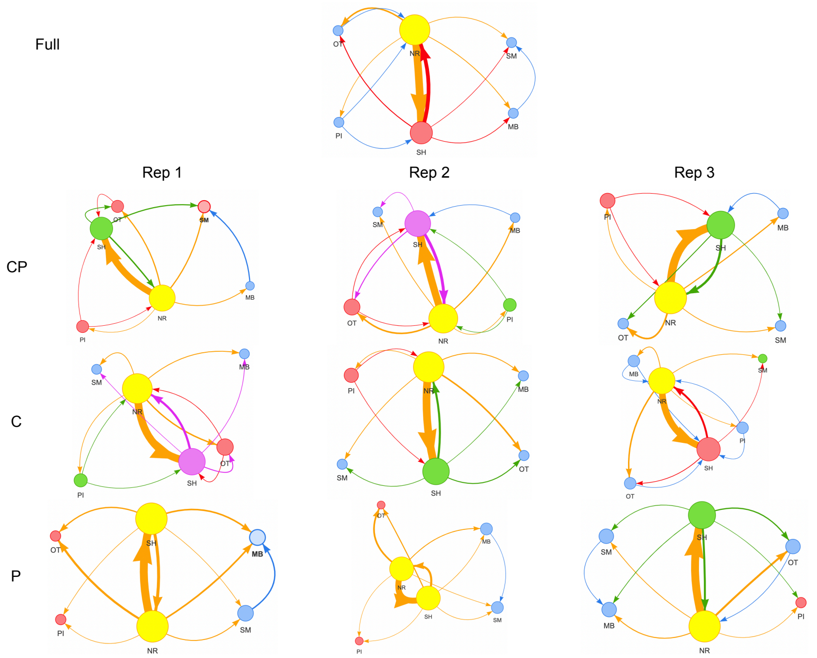


Figure 5: Reconstruction of HIV-1 subtype C transmission networks among PGRHA using the *full* and subsampled datasets visualized using StrainHub v.1.1.2. Nodes are scaled based on degree centrality, where larger nodes correspond to larger centrality.

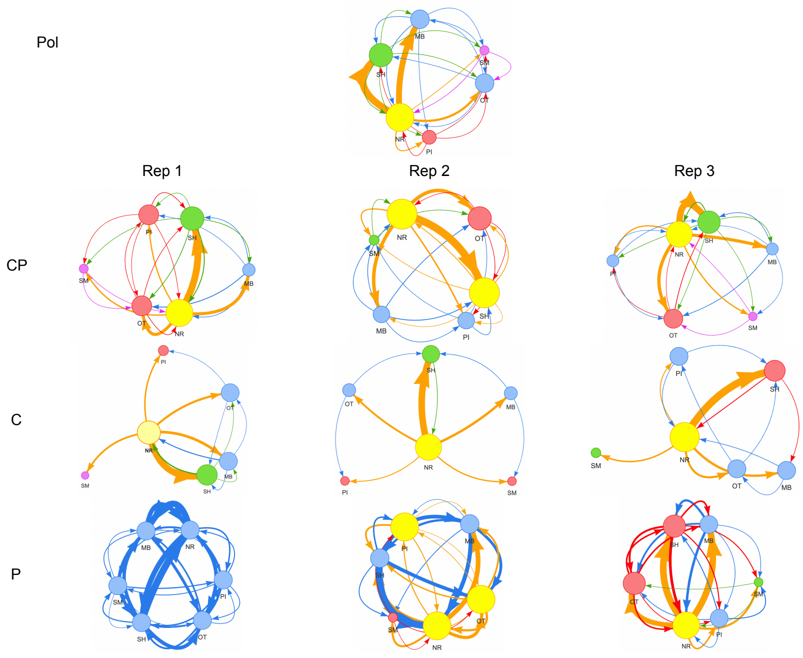


Figure 6: Reconstruction of HIV-1 subtype C transmission networks among PGRHA using the *pol* and subsampled datasets visualized using StrainHub v.1.1.2. Nodes are scaled based on degree centrality, where larger nodes correspond to larger centrality.
